# AMH regulates ovary size by counteracting ovarian follicle cluster effects

**DOI:** 10.1101/2024.08.15.607694

**Authors:** Carmen Lim, Lynn Yiew, Nicholas J. Anderson, Peter Smith, Laurel Quirke, Alan Carne, Urooza Sarma, Ryan Rose, Martha Nicholson, Christine L. Jasoni, Ping Liu, Simone Petrich, Jenny Juengel, Michael W. Pankhurst

## Abstract

Serum anti-Müllerian hormone (AMH) is the primary clinical indicator of mature oocyte counts in the ovaries, but its biological role remains poorly understood. Mammalian ovaries have a finite lifetime of oocytes that are slowly depleted as the dormant follicles housing the oocytes initiate maturation. Less than 0.1% of these follicles will reach maturity and ovulate an oocyte. Recent studies suggest that AMH is a key regulator that removes most of these follicles at early stages of follicle maturation. Most AMH is secreted as an inactive precursor protein, and we show that the required activating-enzymes are largely present outside the follicle. We then measured AMH concentrations in ovarian stroma with microdialysis showing that activity is confined to a short range from the site of secretion. To examine short-range interactions between follicles, we reconstructed the ovarian follicle positions from sheep ovaries in 3D space. This showed that most early follicles develop in proximity to more advanced follicles. Active immunisation of sheep against AMH to inhibit signalling, greatly expanded early follicle numbers, but almost entirely in proximity to large follicles. Large follicle proximity appears to greatly enhance early follicle survival, and AMH appears to attenuate this effect to prevent follicle overgrowth beyond sustainable limits.

## Introduction

Anti-Müllerian hormone is a TGFβ superfamily member first discovered as the fetal testicular hormone that removes the female reproductive tract (paramesonephric/Müllerian duct) from male embryos^1,2^. It was considered a male hormone for decades until AMH expression was observed in the adult ovary^3^ but its ovarian function remains unclear. The ovary is bestowed at birth, with a non-renewable supply of dormant (primordial) follicles that house the immature oocytes. Across the reproductive lifespan, primordial follicles are constantly activated to ensure a continuous supply of developing follicles is available for ovulation each ovarian cycle. As soon as a primordial follicle activates and begins development towards full maturity, AMH is expressed in the granulosa cells surrounding the oocyte^4^. AMH expression increases throughout primary, preantral and antral follicle development until the final days of folliculogenesis when the follicle prepares for ovulation and AMH synthesis tapers off^5^.

Most ovarian hormones have clearly defined roles; estrogens are the primary sex hormones and orchestrate ovulation, progesterone facilitates pregnancy, inhibins regulate gonadotropin secretion, but a similar summary statement does not yet exist for ovarian AMH. AMH circulates in blood but the main site of AMH receptor (AMHR2) expression in adults is the granulosa cells^3,4^, suggesting that AMH might be primarily an autocrine signal, rather than a hormone. Paracrine AMH secreted from developing follicles is known to inhibit primordial follicle activation^6^, as the primordial follicles express AMH receptors but not AMH itself^7^. This function has been proposed to prolong the length of the female reproductive lifespan but *Amh*^−/−^ mice do not have reduced lifetime fertility and only have minor reductions in the timing of menopausal onset^8^. AMH also induces atresia (follicle degeneration) in early (preantral) follicle development^9,10^ and reduces the production of estrogens in nearly mature (antral) follicles^11^. Antral follicles require follicle-stimulating hormone (FSH) to grow and survive^12^ and some studies suggest that AMH inhibits this activity^13,14^, possibly to prevent follicle maturation from progressing too quickly^5^. However, *Amh*^−/−^ mice ovulate functional oocytes from apparently normal mature follicles^15,16^, suggesting that this is also not an essential role of AMH.

Perhaps the most confusing aspect of AMH signalling in the ovary is that it initially inhibits dormant primordial follicle activation^6^, then causes preantral follicle atresia^9,10^, before switching back to trophic support as follicles transition from preantral to small antral and finally, becoming growth-inhibiting in large antral, and fully mature follicles^11,17,18^. We hypothesise that this allows small antral follicles to produce large quantities of AMH without ill-effect, and then secrete that AMH outside the follicle to cause atresia on nearby preantral follicles. Most AMH is secreted into follicular fluid as an inactive proprotein precursor (proAMH)^19^ but at some stage before the AMH leaves the ovary and enters circulation, it is cleaved to the active form^20,21^. This study aimed to determine the site of AMH activity in the ovary by examining locations of activity for the enzymes that cleave and activate proAMH, subtilisin/kexin type proprotein convertases (PCSKs)^22^. Finally, we aimed to examine 3D spatial interactions between the large follicles that produce the bulk of the AMH and the nearby preantral follicles that are susceptible to AMH-mediated atresia.

## Materials and Methods

### Participants

Human ovary tissue was sourced from patients with *BRACA1*, *BRACA2* or *RAB51* mutations undergoing prophylactic oophorectomy at Dunedin Hospital, Dunedin, New Zealand. Five patients were recruited between July 2019 and February 2021. Patients were eligible for recruitment if they were >18 years of age, premenopausal and had an indication for removal of one or both ovaries. Exclusion criteria were pregnancy, lactation, reproductive endocrine disorders or endometriosis affecting the ovaries. Ovaries were transported immediately from the surgical theatre to the research laboratory for microdialysis measurements and sampling of ovarian follicular fluid. Microdialysis sampling was limited to 1 hour to enable the ovaries to be transported to the pathology laboratory for fixation within 3 hours of surgery. The sampling procedures had no effect on the quality of the subsequent histology and pathology assessments. No patients were found to have malignancies upon pathology examination. Ethical approval for this experiment was granted by the Heath and Disabilities Ethics Committee, New Zealand Ministry of Health. All participants provided written informed consent.

### Animals

All studies comparing *Amh*^+/+^ and *Amh*^−/−^ mouse (Behringer et al., 1994) ovary tissues utilised mice on a C57Bl6/J background. Adult mice were between 42 and 120 days of age and neonates were used at postnatal day 2, where day zero was the day of birth. The animals were housed in a climate-controlled environment, with 12:12 hour light/dark cycles and free access to water and standard rodent chow.

Sheep used in the AMH-immunisation experiments were Romney breed aged between 2-5 years and all had given birth in at least one prior season. Sheep ovaries used in the microdialysis experiments were obtained from a local abattoir and were therefore, from mixed breeds.

All experiments on live animals or tissue from animals bred specifically for experimentation (i.e. not obtained from an abattoir) were either approved by the University of Otago Animal Ethics Committee or the AgResearch Animal Ethics Committee.

### Microdialysis

Sheep ovaries were obtained from the Silverfern Farms Finegand Abbatoir, Balclutha, New Zealand and were isolated immediately after the sheep had been slaughtered. The ovaries were transported back to the laboratory in Dunedin within 1 hour, where they immediately underwent microdialysis. Human ovaries were obtained for the procedure, as described above. During microdialysis ovaries were contained under an immersion buffer (0.9% NaCl, 0.01 mol/L HEPES, 1% BSA, pH 7.4).

AtmosLM™ microdialysis probes with a 1000 kDa molecular weight cut-off pore-size (Eicom, Cat# PEP-12-04) were inserted into ovarian stroma adjacent to antral follicles estimated to be 2-5 mm in diameter. A 19-guage guide needle was first inserted with the microdialysis probe then lowered down through the needle. Perfusion buffer (0.9% NaCl, 0.01 mol/L HEPES, 10%w/v BSA, pH 7.4) was pumped into the probe via a syringe pump and from probe to collection vessel via a peristaltic pump, with flow rate set to 2 µL/min. Microdialysate was collected over 1 hour. At the end of the procedure, follicular fluid samples were taken using a 29-guage needle. Follicular fluid was centrifuged at 6700 ×g for 5 minutes to pellet out granulosa cells, then the supernatant was transferred to a clean tube and was frozen and stored at −80°C.

Microdialysis of recombinant human proAMH determined that the transfer efficiency into the microdialysate was 1.7%. AMH levels were assayed by ELISA (sheep: AMH (Ovine) ELISA, AnshLabs, Cat# AL-155, human: Ultra-Sensitive AMH, AnshLabs, Cat# AL-105) then the assayed values were adjusted by a factor of 58.8 to determine the stromal interstitial fluid AMH concentrations. The detection limit for AMH was 3.3 pmol/L in human follicular fluid and 10.5 pmol/L in sheep follicular fluid, after correction for extraction efficiency and sample dilution. Follicular fluid samples were diluted in assay buffer before AMH quantification by ELISA and correction for dilution factor.

### Protease-activated fluorogenic reagent imaging

Mouse ovaries from C57Bl6/J mice were dissected immediately after euthanasia. The ovaries were cut into fragments and were incubated in phenolred-free DMEM/F12 medium (ThermoFisher, Cat# 11039021) containing 6 µmol/L Rh110 furin substrate (Sensolyte® Rh110 Furin Activity Assay kit, Anaspec, Cat# AS-72256) and 1.6 mmol/L Hoechst 33342 DNA label (ThermoFisher, Cat# H3570) at 37 °C, 5% CO_2_ for 2 hours with gentle agitation every 30 minutes. The reagent is non-fluorescent but protease activity liberates fluorescent rhodamine 110. Rhodamine 110 accumulation of the incubated ovary fragments was quantified by confocal microscopy imaging with 488 nm excitation, 555 nm emission. Hoechst 33342 was imaged under 405 nm excitation, 450 nm emission. Fluorescence intensity was quantified using FIJI software using the ‘plot profile’ function. Pixel intensity of rhodamine 110 fluorescence was determined along straight-line traces intersecting with the centre of small-medium antral follicles. The angle of the line was predetermined by a random number generator. The mean pixel intensity of each zone was combined (µ_1_ = antral space, µ_2_ = granulosa, µ_3_ = theca, µ_4_ = stroma), and calculate the total pixel intensity within the trace; µ_T_ = (µ_1_ + µ_2_ + µ_3_ + µ_4_)/4. For each follicle (n = 9), the mean pixel intensity of each zone was divided by µ_T_ to calculate a relative fluorescence unit (RFU) value. Repeated-measures ANOVA with Tukey’s B post-hoc test was used to investigate differences in fluorescence across the anatomical zones.

### Neonatal ovary cultures

Neonatal mice were euthanised by decapitation and the ovaries were dissected from the bursa and were transferred into 2 µL media droplet on a transwell insert (0.4μm pore size, Millicell, Merck, Cat# PIHP01250) floating on 0.5 mL of DMEM/F-12 media, 100 U/mL penicillin/streptomycin (Gibco, Cat# 1520096), ITS-G (Gibco, Cat# 41400045) in a 24-well plate at 37°C, 5% CO_2_. Cultures were treated with varied concentrations of recombinant human AMH_C_ (R&D systems, cat# 17-37-MS-010) and were fixed in Bouins fixative for haematoxylin and eosin histological staining and follicle counting.

### Neonatal ovary-adult follicle co-cultures

Neonatal ovaries were co-cultured with adult follicles (100-150 µm diameter) dissected from the ovaries of adult *Amh*^+/+^ and *Amh*^−/−^ mice with 30-guage needles. Follicles were placed adjacent to neonatal ovaries from *Amh*^+/+^ mouse pups and were cultured for 72 hours. The cultures were observed daily and all adult follicles were confirmed to have become fused with the neonatal ovary within 24 hours. The tissues were fixed in 4% paraformaldehyde and were blocked with 5% normal donkey serum (NDS) (Merck, Cat# D9663, RRID:AB_2810235), 0.2% tween-20 in phosphate buffered saline (PBS) for 3 hours. Primary antibodies were 1 μg/mL mouse anti-DDX4 (Abcam, Cat# ab27591, RRID:AB_11139638), 0.05 μg/mL goat anti-AMH (MIS C-20, Santa Cruz Biotechnology, Cat# SC-6886, RRID:AB_649207) applied for 5 days at 4°C. Secondary antibodies were 1 μg/mL donkey anti-mouse IgG-DyLight 488 fluorophore (ThermoFisher Scientific, Cat# SA5-10166, RRID:AB_2556746), and 1 μg/mL donkey anti-goat IgG-DyLight 594 (ThermoFisher Scientific, Cat# SA5-10088, RRID:AB_2556668) and 0.3 μmol/L of 4’, 6-diamidino-2-phenyllindole (DAPI) was included in the solution for a 4 day incubation at 4°C. Z-stacks were obtained by confocal microscopy. Note that the DAPI fluorescence channel has undergone a digital linear-adjustment to increase the brightness to a visible level in the displayed images.

### Follicle culture

A previously published method for follicle culture was adapted for these experiments^23^. One follicle per well was cultured in round-bottom 96-well plates containing 100 µL of MEMα media with 100 U/mL penicillin/streptomycin (Gibco, Cat# 1520096), ITS-G (Gibco, Cat# 41400045) and 355 mIU/mL FSH (Genway, Cat# GWB-3EE59E). Follicles were isolated from the ovary with 30-gauge needles with some theca and stroma still attached and were cultured for 96 hours at 37°C, 5% CO_2_. Follicles were imaged for diameter measurement daily and 50% of the media in the well was replaced with fresh media once each day. Follicles were excluded from the analysis if there was a decrease in follicle size, loss of spherical morphology or extrusion of the oocyte from the granulosa cell layer.

### Statistical Analysis

Group differences between protease-activated fluorogenic signals across ovarian regions were compared with 2-tailed, 1-way ANOVA with Tukey B posthoc test. Group differences between primordial-primary follicle ratios in neonatal ovary-adult follicle co-cultures were compared with 2-tailed, 2-way ANOVA with Tukey B posthoc test. *Amh*^+/+^ and *Amh*^−/−^ follicle growth rates were compared with 2-tailed, 2-way repeated-measures ANOVA with LSD posthoc test. Differences in sheep ovary follicle counts were compared by 2-tailed Student t-test. Kolmogorov-Smirnov tests were used to examine whether distributions of distance between large follicles and 1-layer or 100-200 µm follicle were significantly different. The analysis was conducted within each treatment group for each of the large follicle size classes. Analyses were performed with SPSS v25 (IBM corporation) or Prism 5 (Graphpad software).

Descriptions of AMH protein isoform expression in assisted reproduction patients, western blotting/2D PAGE isoelectric focusing, procedures for active AMH immunisation in sheep and 3D follicle mapping are explained in the supplemental methods.

## Results

### AMH-signalling activity is strongest adjacent to the secreting-follicle

Human antral fluid AMH concentrations have been reported to be as high as ∼9 nmol/L^5^ and serum concentrations are usually below ∼50 pmol/L^24^, but stromal AMH concentrations have not been quantified. Stroma concentrations were determined in sheep ovaries using microdialysis ranging from 90-410 pmol/L (Fig. 1A). This indicates that AMH becomes rapidly diluted as it diffuses out in ovarian stroma. The finding was also confirmed in human ovaries obtained after prophylactic oophorectomy (patient with cancer-causing *BRACA1/2* or *RAD51* mutations). Antral fluid AMH concentrations were very low in some patients, which is a feature of reproductive ageing^25^. However, when follicular fluid levels were high, AMH could also be detected in the adjacent stroma, again with substantial dilution.

**Figure 1.**
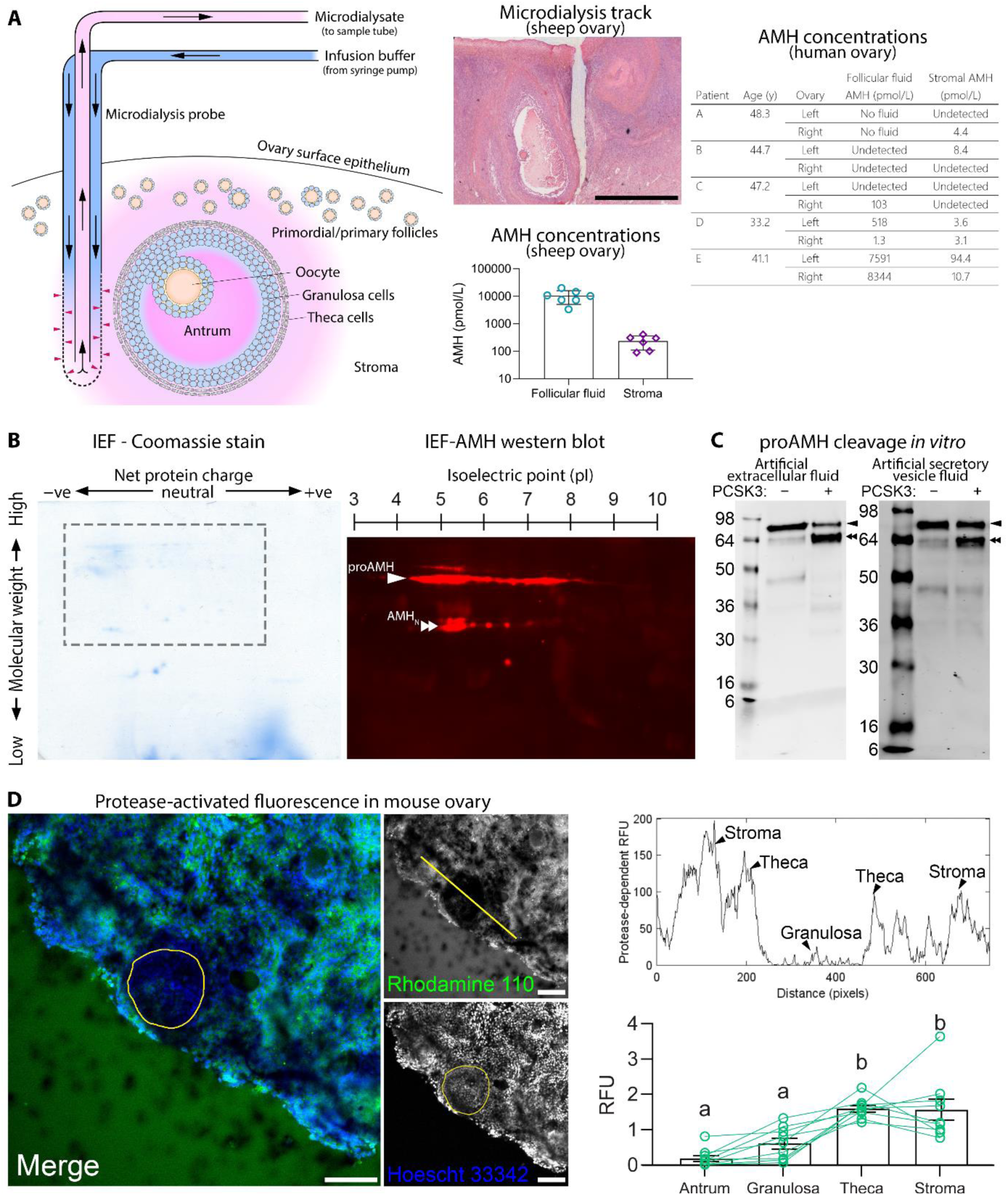
AMH diffusion and proteolytic activation of proAMH in ovarian stroma. (A) Microdialysis probes were inserted adjacent into ovary stroma where dialysis buffer was pumped through the porous probe allowing molecules up to 1000 kDa in molecular weight to enter the dialysate. The dialysate was collected for AMH assay. Needles were placed adjacent to an antral follicle as shown in the hematoxylin and eosin-stained sheep ovary section (Scale bar = 1 mm). AMH concentrations in follicular fluid or adjacent stroma were quantified in sheep and human ovaries. (B) 2D-PAGE separating recombinant AMH protein by size on the vertical dimension and isoelectric focusing (IEF) on the horizontal direction. Western blot was conducted on the dashed area using an antibody against the AMH N-terminal fragment (AMH_N_). The recombinant AMH preparation contains predominantly proAMH (arrow) and lesser quantities of AMH_N_. (C) Western blot of recombinant proAMH treated with (+) or without (−) recombinant PCSK3 in artificial extracellular fluid or artificial secretory vesicle fluid. ProAMH appears as a 72 kDa band (arrowhead) and the cleaved AMH_N_ fragment as a 64 kDa band (double arrowhead). (D) PCSK activity in mouse ovary fragments was visualised with a fluorogenic reagent that releases rhodamine 110 when exposed to protease activity (green: rhodamine 110 and blue: Hoescht 33342 DNA-stain, scale bar = 100 µm). A yellow ring denotes the basal lamina that separates the granulosa and theca layers. The upper graph shows the relative fluorescence intensity units (RFU) along the straight-line trace shown in the rhodamine 110 image and the lower graph shows mean (±SE) fluorescence intensity in the antrum, granulosa, theca and stroma was compared with 1-way ANOVA (p < 0.001) and Tukey’s B post-hoc test (regions that share the same letter were not significantly different, n = 9).

It remains unclear why most of the AMH in follicular fluid is uncleaved proAMH^19^ but the site of extracellular cleavage could reveal information about the site of AMH action. We incubated recombinant proAMH with its cleavage enzyme, PCSK3 in an artificial extracellular fluid, leading to efficient cleavage (Fig. 1C). Cleavage was less efficient when the PCSK3 incubation was conducted in artificial Golgi apparatus/secretory vesicle fluid (Fig. 1C), suggesting that AMH is protected from intracellular cleavage and predisposed for extracellular cleavage. To determine the potential sites of proAMH cleavage, *ex vivo* mouse ovary fragments were incubated with a fluorogenic regent that releases rhodamine 110 when exposed to PCSKs. The liberation of fluorescence was low in antral fluid and the granulosa layer but was high in stroma and theca (Fig. 1D). This observation is consistent with the conversion of proAMH to AMH_N,C_ in the theca and stroma and explains how AMH_N,C_ is present in circulation (theca is vascularised, granulosa is avascular).

The granulosa cells of ovarian follicles form a molecular sieve that allows small molecules (<100 kDa) to pass freely, but retains large molecules (>500 kDa) in the antrum to encourage osmotic fluid-retention^26^. Dimeric AMH has a molecular weight of 140 kDa and intermediate-size proteins (100∼150 kDa) can pass freely through the sieve if they carry a positive charge, but are restricted if they have a negative charge^27^. Isoelectric focusing to determine the net charge of AMH (Fig. 1B, Supplemental Fig. 2) showed variable protein charges, indicative of multiple post-translational modification variants. Densitometry indicated that 66% of the AMH was negatively charged, 18% was neutral and 16% was positive. It is not clear why a large proportion of AMH carries a charge that would slow its egress from the follicle, but it may be a mechanism to prevent freshly generated AMH_N,C_ in the theca/stroma from diffusing back into the granulosa layer. Collectively, these experiments suggest that the target site of AMH activity is either the theca layer or the ovarian stroma immediately adjacent to the AMH secreting-follicle.

Prior evidence suggested that AMH activation is further influenced by luteinising hormone^28,29^. However, we examined the ratios of proAMH and total AMH in the serum of patients undergoing gonadotropin hormone stimulation for IVF and found no change after LH-receptor stimulation (Supplemental Fig. 3). The lack of change in proAMH to total AMH levels indicates that cyclic changes in gonadotropins are unlikely to affect rates of proAMH conversion to AMH_N,C_ across the ovarian cycle.

### Endogenous AMH has limited short-term effects on primordial follicles

In AMHR2-transfected transformed cell lines, the dynamic range of AMH activity occurs from 0.1∼4 nmol/L ^30,31^. We treated neonatal ovary cultures with recombinant AMH and found inhibition of primordial follicle activation occurred over a similar range (Fig. 2A), suggesting that ovarian tissues have the same dose-response curve. We then investigated whether endogenous AMH released from developing follicles was sufficient to induce the same effect. A single developing follicle derived from adult *Amh*^+/+^ or *Amh*^−/−^ mice was fused to an *Amh*^+/+^ neonatal ovary organ culture (Fig. 2B-E). Interestingly, the presence of an antral follicle was sufficient to significantly reduce primordial follicle activation rates but the effect occurred with antral follicle from either *Amh*^+/+^ or *Amh*^−/−^ mice (Fig. 2F). In this experiment, AMH did not have a strong influence beyond the other factors from antral follicles that influence primordial follicle activation (Fig. 2F), in contrast to the strong effects obtained with recombinant AMH (Fig. 2A).

**Figure 2.**
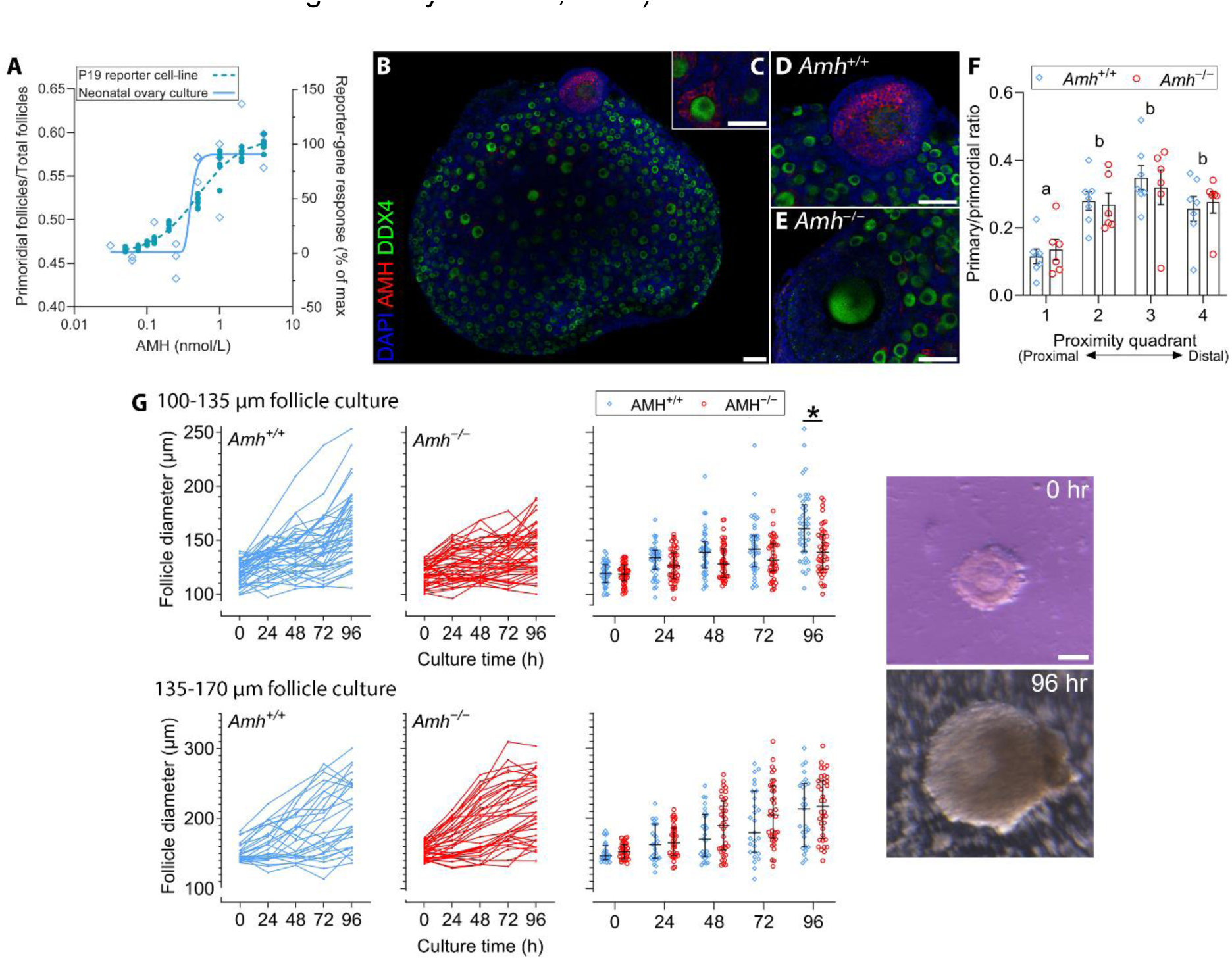
The effects of AMH on primordial and antral follicles. (A) Dose-response curve of recombinant AMH on primordial follicle activation rates in neonatal mouse ovary cultures (blue open diamonds) and AMHR2-transfected P19 reporter cell line (green closed circles). Activation is expressed as the number of remaining primordial follicles as a fraction of all follicles (primordial + developing). (B) Confocal microscope images of antral follicles from adult *Amh*^+/+^ or *Amh*^−/−^ mice fused to *Amh*^+/+^ neonatal ovaries in co-culture with immunofluorescence for AMH, DDX4 (oocyte marker) and DAPI (DNA label). (C) Primary follicle expressing AMH in granulosa cells, (D) adult follicle from an *Amh*^+/+^ mouse and (E) adult follicle from an *Amh*^−/−^ mouse, scale bars = 50 µm. (F) Distances between the antral follicle and all primordial or primary follicles in the antral follicle-neonatal ovary co-culture were divided into quadrants to investigate the primordial follicle activation rate at increasing distances from the antral follicle (2-way ANOVA with Tukey’s B post-hoc test. Groups that share letters are not significantly different). (G) *Amh*^+/+^ and *Amh*^−/−^ small antral follicle growth in culture (* indicates p<0.05, repeated measures ANOVA with Tukey’s B post-hoc test) with representative images of follicles at 0 and 96 hours in vitro.

### Endogenous AMH has limited short-term effects on small antral follicles

The effects of recombinant AMH on early antral follicle development have been contradictory, with both growth-promoting and growth-inhibitory effects observed, ever since the initial experiments^13,32^. The rapid drop in stromal AMH levels within a short distance of the follicle suggests that nearly all AMH found in a follicle is produced endogenously. We investigated the effects of endogenously produced AMH in late preantral or early antral follicles dissected from adult *Amh*^+/+^ or *Amh*^−/−^ mice (Fig. 2D). When late preantral follicles (100-135 µm diameter) were cultured for 24 hours, *Amh*^+/+^ follicles grew slightly larger than *Amh*^−/−^ follicles (*p*<0.001). However, there was no difference in growth-rates when early antral follicles (135-170 µm diameter) from *Amh*^+/+^ and *Amh*^−/−^ mice were cultured in the same manner. These relatively modest effects of AMH on primordial and late preantral/early antral follicles suggest that AMH might be part of a milieu of growth factors influencing follicle function but is not a strong regulator by itself.

### Antral follicle AMH induces atresia in immediately-adjacent preantral follicles

In our studies in mice, the most consistent effect of AMH that we have observed is the induction of atresia in early preantral follicles^9,10^. We investigated this effect in a sheep model where animals were actively immunised against AMH protein or a vehicle-carrier protein, as a control. After 6 months of injections, the AMH-immunised sheep ovaries had no significant reductions in the smallest preantral follicles with 1, 2 or 3 layers of granulosa cells (Fig. 3D). However, counts of the largest (100-200 µm diameter) preantral follicles were significantly expanded in the AMH-immunised sheep compared to control ovaries. This difference persisted in the early antral follicles (0.2-2 mm diameter) but was lost at the larger antral stages, which is very similar to the phenotype we observed in *Amh*^−/−^ mice^10^. There were also no differences in mean ovulation rates between the AMH-immunised (1.68 ovulations per cycle), and control (1.68 ovulations per cycle) sheep.

**Figure 3.**
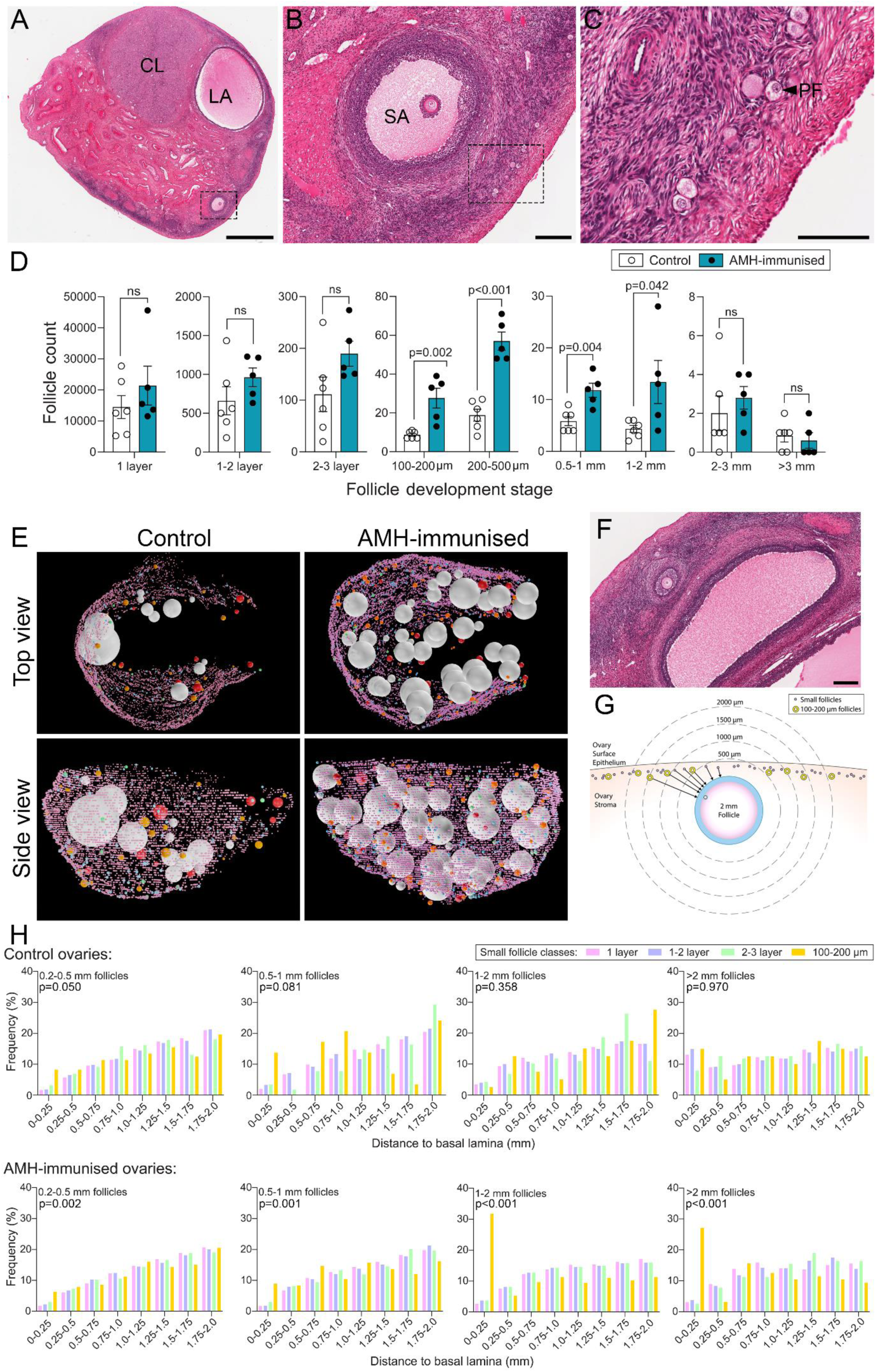
Three-dimensional reconstruction of sheep ovary follicles. All data shown were collected from half of a bisected ovary. (A-C) Haematoxylin and eosin-stained sheep ovary sections (abbreviations: CL; corpus luteum, LA; large antral follicle, SA; small antral follicle, PF; primordial follicle). The dashed line in panels A and B shows the region depicted in the subsequent panel. Scale bars: 2mm, 200 µm and 100 µm in A, B and C, respectively. (D) Follicle counts in each ovary with small follicles classified by the number of granulosa cell layers (1 layer, 1-2 layers or 2-3 layers) and large follicles classified by follicle diameter (p-values determined by Student’s t-test). (E) Follicle locations in 3D space in control and AMH-immunised sheep ovaries. (F) Example of a 100-200 µm preantral follicle in proximity to an antral follicle in an AMH-immunised sheep ovary. Scale bar: 200 µm. (G) Schematic depicting follicle distance measurements between preantral and antral follicles. (H) Distributions of distances between large antral follicles and 1 layer, 1-2 layers or 2-3 layer or 100-200 µm follicles within 2000 µm of the antral follicle basal lamina. Data are combined across all antral follicles from 5 control sheep ovaries or 4 AMH-immunised sheep ovaries. P-values were generated with Kolmogorov-Smirnov tests between the 1-layer and 100-200 µm follicles.

Unlike mouse ovaries which are densely populated with follicles, the follicles in sheep ovaries were more dispersed around the ovary. It was therefore unclear how the removal of AMH signalling by immunisation could affect preantral follicles across the whole ovary, when stroma concentrations suggest it can only act within a short distance of the follicle it is secreted from. To investigate this further, we mapped the 3D coordinates of 29,212 follicles from 4 AMH-immunised sheep ovaries and 29,322 follicles in 6 control ovaries (Fig. 3E, Supplemental fig. 4&5, Supplemental movie). In control sheep, the follicles with 1-layer of granulosa cells (presumptive primordial follicle), 1-2 layers or 2-3 layers were found distributed at all locations within the ovarian cortex. However, the 100-200 µm follicles were often found in proximity to larger follicles or in clusters with other similarly-sized follicles suggesting proximity to other follicles highly increases late preantral follicle survival rates.

To determine if increased preantral follicle survival in AMH-immunised sheep was based on proximity to antral follicles, we calculated the distance between small follicles and large follicles and plotted the frequency distributions (Fig. 3H). Almost all of the increase in 100-200 µm preantral follicles could be found within 250 µm of the edge of a follicle larger than 1 mm in diameter (see example, Fig. 3F). This indicates that AMH inhibits the excessive preantral follicle survival that occurs in close proximity to larger follicles.

## Discussion

In 2009, Da Silva-Buttkus, et al., published an article predicting the existence of an unidentified local inhibitor of follicle growth in postnatal ovaries^33^. Here we use 3D spatial analysis to show that AMH is one such local inhibitor, and that these effects occur in adult ovaries, in large animals. Furthermore, this finding suggests that AMH action is required to balance the growth-promoting and survival-enhancing effects that antral follicles impart to nearby preantral follicles.

The ovaries of the average 20-25-year-old woman will activate ∼450 primordial follicles per month^34^ and if all were allowed to reach 4 mm diameter (medium antral stage), their combined volume would be 15.1 cm^3^, which is larger than the average volume of both ovaries combined (∼9 cm^3^)^35^. It takes an estimated 3 weeks for a follicle to grow from early-antral (2 mm), to the fully-mature preovulatory size (20 mm)^36^ and numerous follicles will attempt this each cycle. Therefore, the ovarian stroma needs to repeatedly accommodate large structural changes and presumably, the capacity of the ovary for cellular restructuring has an upper limit. This is evidenced by the accumulation of connective tissue in the ovary with ageing^37^ and one theory posits that elevated rates of ovarian mitosis may contribute to oncogenesis^38^. Removing the majority of excess follicles at the preantral stage may have evolved to mitigate the high metabolic, or mitotic burden caused by larger antral follicles.

Follicles produce a range of growth factors that likely support follicular growth and survival in an autocrine manner^39^. However, these same growth factors diffusing out into stroma may also act on nearby follicles and may explain why developing follicles tend to be found in clusters. This is supported by studies showing that paracrine factors from healthy ovary fragment transplants can rescue follicle development in ovaries rendered non-functional by chemotherapy^40^. Antral follicles also promote angiogenesis in their thecal layer to increase the delivery of vital nutrients and endocrine signals to the follicle^41^. This may also bring an influx of growth-promoting factors to preantral follicles in the adjacent stroma. If our interpretation is correct, then antral follicles secrete AMH to counterbalance the ‘proximity effect’ that favours preantral follicle survival.

There is increasing evidence that ovarian sites beyond the granulosa cell layer are targets of AMH. Recombinant AMH_C_, the 25 kDa receptor-binding fragment, accumulates in stroma but not granulosa after intraperitoneal injection^42^. This suggests that additional mechanisms exist to retain AMH in ovarian stroma. AMH receptors have recently been detected in theca cells^43^, and AMH has been shown to inhibit theca cell androgen production^44^. This local androgen production appears to be important for preantral follicle growth and survival^45–47^, and its inhibition may be a mechanism of AMH-mediated preantral follicle loss. However, more research will need to be done *in vivo*, taking steps to ensure that experiments are conducted with physiological levels of AMH. Strong effects of recombinant AMH have been observed on all stages of folliculogenesis *in vitro*, but the same effects have not always been clearly apparent *in vivo*.

The present study suggests that the primary role of AMH in the ovary is to prevent excessive numbers of follicles from surviving beyond the preantral stage. An important implication of these findings is that the composition of the follicle population in an ovary in the current cycle can influence the composition and structure of future cycles. More research will be needed to determine how this affects the ovary during ageing, infertility and pathological conditions.

## Acknowledgements

Nicola Batchelor, Andrew McNaughton, Otago Micro and Nanoscale Imaging Unit, Otago Histology Services Unit, and Biomedical Research Facility, University of Otago and AgResearch Farm staff for technical assistance. Danny Gutsell, Ted Dons and staff from the Finegand plant at Silverfern farms, Balclutha, New Zealand for provision of sheep ovaries. Nigel Groome for antibodies used for immunisation titre analysis. Fertility SA for collection, preparation and storage of serum samples. M.W.P. discloses support for the research of this work from Sir Charles Hercus Research Fellowship from the Health Research Council of New Zealand [grant number 18-027].

## Author contributions

C.L.; experimentation/analysis, L.Y.; experimentation/analysis, N.J.A.; experimentation/analysis, P.S.; design/experimentation, L.Q.; experimentation/analysis, A.C.; design/experimentation, U.S.; design/experimentation, R.R.; design, M.N.; design, C.L.J.; design, P.L.; design, S.P.; design, J.J.; design/analysis, M.W.P.; design/experimentation/analysis/manuscript draft. All authors have approved the submitted version and agree to be personally accountable for the work described herein.

## Data Availability statement

The datasets generated during and/or analysed during the current study are available from the corresponding author on reasonable request.

## Supplemental methods

### In vitro fertilisation ovarian stimulation patient recruitment

Serum samples were obtained from 75 patients attending Fertility SA, St Andrews Hospital, Adelaide, Australia, a privately owned ART clinic to measure proAMH and total AMH levels during controlled ovarian stimulation. The mean (SD) age was 36.1 ± 4.8 years (range: 24-45). Standard gonadotrophin releasing hormone antagonist stimulation cycles were used with doses of FSH individualized to each patient, based on ovarian reserve parameters, previous IVF response, or both. The FSH doses ranged from 100 IU to 400 IU. The first serum sample was obtained from patients when three follicles measuring wider than 17 mm were identified using transvaginal ultrasonography, and a HCG or agonist trigger injection was given within 24–48 h. A second blood sample was taken from each participant at the time of oocyte retrieval (36 hours post-trigger injection). Immunoassays for proAMH and total AMH were performed as previously described^1^. Of the 75 participants assay, only 15 had sufficiently high AMH levels for reliable quantification of the ratio of proAMH to total AMH and thus, inclusion in the final analysis (Total AMH >10 pmol/L or proAMH <0.9 pmol/L). The ratio was converted to an AMH prohomone index with the following formula; API = [proAMH]/[total AMH] × 100. Ethical approval for this experiment was granted by The University of Otago Human Ethics Committee (Health) and the St Andrew’s Hospital Human Research Ethics (STAND Project #93), Adelaide, Australia. All participants provided written informed consent.

### Active AMH-immunisation

Total cellular RNA was isolated from ovine ovarian tissue using TRIzol™ (Invitrogen) according to the manufacturer’s instructions. First strand cDNA was synthesised from 5 µg of total RNA using SuperScript™ First Strand Synthesis System for RT-PCR (Invitrogen). The PCR primers (Forward: 5’<u>ggatcc</u>gagcaccggagccgcggctgc and Reverse: 5’g<u>aagctt</u>ccggcagccgcattcggtg) were designed to amplify the nucleotides encoding amino acids 467-575 of the ovine AMH (oAMH) protein (NP_001295528.1). The forward and reverse primers included BamHI and HindIII sites (underlined), respectively, required for subcloning into pQE31 expression vector. The PCR reaction employed the use of the GC-Rich PCR System (Roche). Following denaturation at 94°C for 3 minutes, the cDNA was amplified using 35 cycles of denaturing for 1 minute at 94°C, 1 minute of annealing at 60°C, and extension at 72°C for 2 minutes. A final extension was performed for 10 minutes at 72°C. Amplification products were digested with BamHI and HindIII and subjected to electrophoresis through a 1% (w/v) low melting point agarose gel. Product of the expected size was ligated into expression vector pQE31 (Qiagen) and the identity of the cloned insert confirmed by DNA sequencing. The recombinant plasmid was transformed into *E. coli* strain M15(pREP4) (Qiagen) followed by protein production induced by isopropylthio-b-galactoside and confirmed on western blots by labelling with anti·His antibody (Qiagen) and anti-AMH antibody (kindly provided by Nigel Groome). Induced cells were lysed by lysozyme treatment and sonification. The recombinant protein oAMH was solubilised in 8 mol/L Urea and purified using Ni-NTA-agarose (Qiagen) utilising the pQE31 C-terminal six-histidine His-tag vector sequence. Bound oAMH was eluted in the presence of 8 mol/L urea/250 mmol/L imidazole. Equal amounts of the *E. coli* produced oAMH and the carrier protein keyhole limpet hemocyanin (KLH) were conjugated in the presence of 1% gluteraldehyde and 1 mol/L glycine. Conjugation was confirmed by SDS-PAGE.

Ewes were immunised with 0.4 mg of either KLH (control n = 6) or AMH-KLH (n = 5) in 1 mL of Freund’s complete adjuvant, given 4–5 months before the onset of the breeding season. The ewes then received 5 more injections every 28 days with 0.2 mg of KLH or KLH-AMH in 1 mL of sorbitan triolate, Tween 85, Marcol 52 mineral oil; 1:1:8 v/v/v). After the fifth injection, vasectomized rams with marking harnesses were run with the ewes to monitor estrous cycles with estrous cycle length calculated as the number of days between first and successive markings by the vasectomized ram. Ovulation rates were determined by laparoscopy over 3 cycles with final ovulation rate determined upon tissue collection after euthanasia. In addition, ovulation rate of all ewes was determined by laparoscopy 3 - 4 weeks prior to ovarian collection. Thus, the control ewes underwent laparoscopy two to three times and the treated groups most commonly once, although some were subjected to laparoscopy two to three times also. Ewes were euthanised using a captive bolt and exsanguination 30 days after the final injection. Ovaries were immersion fixed in Bouin’s fixative overnight at room temperature then were wax-embedded for haematoxylin and eosin histological staining of serial sections with 5 µm thickness. Sera was collected from all immunised ewes prior to immunisation, 2 weeks after the booster immunisation, and at time of ovary collection. Antibody titers were determined by qualitative ELISA. ELISA conditions were determined as follows: wells were coated with 100 µg/well of oAMH protein in 0.05 mol/L sodium bicarbonate buffer, pH 9.6; sera were diluted 1:5000; secondary antibody HRP-conjugated rabbit anti-sheep IgG (Invitrogen) was diluted 1:10,000. All AMH-immunised sheep sera contained anti-AMH antibodies 6 weeks after the first injection and at the time of euthanasia.

### Western blot

Recombinant PCSK3 (8 units) (New England Biolabs, Cat# P8077L) was preincubated in 10 µL of 1.2 mmol/L CaCl_2_, 5 mmol MES (brand), pH 6.0 to activate the enzyme. Four pmol of recombinant human proAMH (PX’ Therapeutics) was added either in a buffer resembling the ion content and pH of extracellular fluid (1.8 mmol/L Ca^2+^, 140 mmol/L Na^+^, 4 mmol/L K^+^, 0.05%v/v Triton X-100 (brand), 100 mmol/L HEPES, pH 7.4) or one resembling conditions in the Golgi complex^2,3^ (0.4 mmol/L Ca^2+^, 12 mmol/L Na^+^, 107 mmol/L K^+^, 0.05%v/v Triton X-100 (brand), 100 mmol/L MES, pH 6.5. The final reaction volume was 40 µL and the incubation was conducted for 24 hours at 37 °C. SDS-PAGE was run on 10% Tris-glycine acrylamide gels with 4% stacking gel at 100 volts for 1.5 h using the Xcell Surelock Mini-Cell system (Invitrogen). Proteins were transferred to a 0.4-µm nitrocellulose membrane (Whatman) at 30 volts for 1 h on ice. Blotting membranes were blocked with Odyssey blocking reagent (Licor) for 30 min then were probed with 0.1 µg/ml primary antibody to the N-terminal fragment of AMH (R&D systems, Cat# AF2748, RRID:AB_2226475) then with 1 µg/mL IRDye 680RD donkey anti-goat IgG antibody (Licor, Cat# 926-68074, RRID:AB_10956736). Immunofluorescence was imaged on an Odyssey infrared fluorescence scanner (Licor).

### Isoelectric focusing and 2D-PAGE

Aliquots of recombinant human AMH (625 ng) were combined with 130 µL of an isoelectric focusing (IEF) rehydration buffer [containing 7 mol/L urea, 2 mol/L thiourea, 2%w/v CHAPS, 50 mM DTT, 5 mmol/L TCEP (triscarboxyethylphosphine), to which 25 µL of IEF buffer concentrate (IPG buffer, GE Healthcare, Cytiva) was added per 0.5 mL of rehydration buffer, just prior to use], and the solution used to rehydrate overnight Immobiline DryStrip pH 3-10, 7 cm, linear gradient IEF strips (GE Healthcare). The rehydrated IEF strips were then subjected to electrophoresis in an IEF flat bed electrophoresis system (IPGphor, Pharmacia Biotech) in a ceramic welled plate with the IEF strips submerged under mineral oil, and using a voltage programme from 200 - 8000 V over 10 h, with a V/h target of 30,000 V/h. After IEF, the strips were applied to an in-house prepared second dimension SDS-PAGE (12.5% acrylamide resolving gel, no stacking gel) and held in place using molten 0.5%w/v agarose. After electrophresis the SDS-PAGE gels were stained overnight with SimplyBlue SafeStain (Invitrogen) and then destained in water. An image of the destained gel was captured using a Canon CanoScan LiDE 600F scanner.

A second identical 2D SDS-PAGE gel underwent electroblotting to transfer the protein components to nitrocellulose membrane in transfer buffer (25 mmol/L Tris, 192 mmol/L glycine, 10%v/v methanol) at 300 mA for 3 hours. Immunolabelling of the 2D blot was performed as described, above. Densitometry measurements were performed using FIJI software (NIH)^4,5^.

### Sheep ovary 3D follicle mapping

Sheep ovaries were bisected in half before embedding for histological sectioning. The whole ovary cross-section of every 20^th^ section (100 µm section interval) was scanned at 20x magnification (Aperio CS2 Digital Slide Scanning System, Leica) and stored as .svs files. Follicles were sub-classified into 1-layer (no more than 1 layer of granulosa cells at any point around the follicle), 1-2 layers (a second layer of granulosa cells starting to form but not completely enveloping the follicle) or 2-3 layers (a complete second layer of granulosa cells present but the third layer remaining incomplete). 1 layer and 1-2 layer follicles were excluded if the cytoplasm was not visible and 2-3 layer follicles were excluded if the nucleus was not visible in the cross-section. The x,y coordinates from the centre of each follicle cross-section was recorded using the notation function in the ImageScope software (v12.4.3, Leica Biosystems Pathology Imaging). Coordinates obtained within individual images were designated x_u_,y_u_ and were recorded in a database. These data were also used to generate follicle counts, with an Abercrombie correction performed to account for differences in follicle diameter^6^.

Due to memory constraints, the .svs files could not be compiled into a Z-stack. Each .svs file had its resolution reduced to 72 dpi and was converted to a .tif file. The re-sized .tif files were loaded into Z-stacks in FIJI using the trackEM automated registration function to align the images correctly in 3D space. The cartesian coordinates of the mid-point and diameter all follicles larger than 100 µm in diameter (> 3 layers) was determined and recorded in the stack. Z-coordinates were determined by multiplying the section number by 100 µm. These coordinates were stored in a separate database where the coordinates were designated x_3_,y_3_,z_3_. Large follicles were classed into specific size classes including 100-200 µm, 200-500 µm, 500-1000 µm, 1000-2000 µm, 2000-3000 µm, and >3000 µm. Follicles were recorded as atretic if the granulosa cells in the mural layer showed extensive pyknosis consistent with apoptosis and/or cellular degeneration of the oocyte. Atresia was not recorded in follicles with ≤3 layers.

The x_u_,y_u_ coordinates are accurate within each image but had not been aligned to the whole ovary x_a_,y_a_,z_a_ coordinates that were generated by the Z-stacks of the resized images. However the re-sized images did not have sufficient resolution to identify preantral follicles. To resolve this problem, the x_u_,y_u_ coordinates generated from the .svs files were mathematically rotated to generate an intermediate x,y coordinate, which was then were translated to align with the x_a_,y_a_ coordinates generated in the fully-aligned low-resolution image Z-stack. The top left and top right corners of each image were designated P1 and P2 respectively (Supplemental fig. 1). P3 was generated from the x-value from P2 and the y-value from P1 to generate a right-angle triangle between the 3 points. The angle, θ, at P1 was calculated using trigonometry: cosθ=base length/hypotenuse length, where, hypotenuse (distance between P1 and P2) and base (distance between P1 and P3) lengths were calculated using the formula for Euclidian distance:

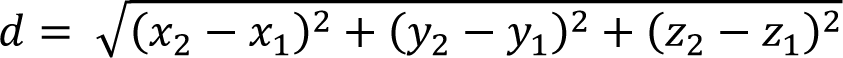

Within the .svs files, P1 represents the origin of the original image but within the Z-stack, P1 has been shifted by the alignment process. The x,y coordinates of P1 indicate how much the image needs to be translated, as this indicates how far the image has been moved from the origin. Therefore, the rotation and translation for each image in the aligned, re-sized .tif Z-stack was determined.

Each preantral follicle x_u_,y_u_ coordinate was rotated by angle θ to calculate the x_r_,y_r_ coordinates. The x,y coordinate of P1 was used to determine the translation distance to generate the x_a_,y_a_ coordinates (Supplemental fig. 1). The z_a_ coordinate was determined by the section number multiplied by section thickness (5 µm). The following formula was used for clockwise rotation and translation; x_a_ = (x_u_cosθ – y_u_sinθ) + x_P1_ and y_a_ = (–y_u_sinθ + y_u_cosθ) + y_P1_. An alternate formula was used for counterclockwise rotation and translation; x_a_ = (x_u_cosθ – y_u_sinθ) + x_P1_ and y_a_ = (y_u_sinθ + y_u_cosθ) + y_P1_.

Once all preantral follicle coordinates had been aligned with the coordinates recorded for large follicles, the data was rendered in 3D using Blender software v3.6 (The Blender Foundation). Data were imported with the Spreadsheet Importer Addon^7^. All follicles larger than 100 µm in diameter were generated with their true diameter. The 1 layer, 1-2 layer and 2-3 layer follicles were assigned slightly larger diameters (60 µm, 120 µm and 160 µm, respectively) to improve their visibility in the rendered images and movies.

Distances between follicles were calculated using Euclidian geometry. The closest distance between the basal laminae of two follicles was calculated using the following formula:

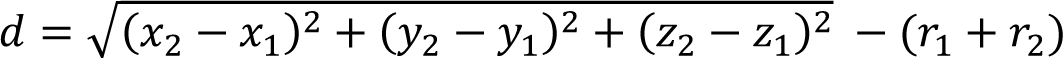

where, *d* is distance, each set of x_a_,y_a_,z_a_ coordinates represents the midpoint of one follicles and *r* and is the radius of each follicle, from midpoint to basal lamina. The radius was set to zero for 1 layer, 1-2 layer and 2-3 layer follicles, due to their small size and the difficulty in obtaining an accurate diameter measurement for individual follicles.

The distances between all follicles in the smaller classes (<200 µm) were determined relative to follicles in the larger classes (>200 µm). In total, 58,534 follicles from the control and AMH-immunised follicles were analysed. The analysis of small follicles (1 layer, 1-2 layer, 2-3 layer or 100-200 µm follicles) within proximity of large follicles was limited to a zone reaching 2000 µm from the basal lamina of the large follicle. A maximum of 3000 data points were randomly selected from each follicle class in each ovary and pooled into control and AMH-immunised ovary groups for analysis. The data were analysed as a frequency histogram to examine which classes of small follicles could exist in proximity to larger follicles.

## Supplemental figure legends

**Supplemental figure 1.**
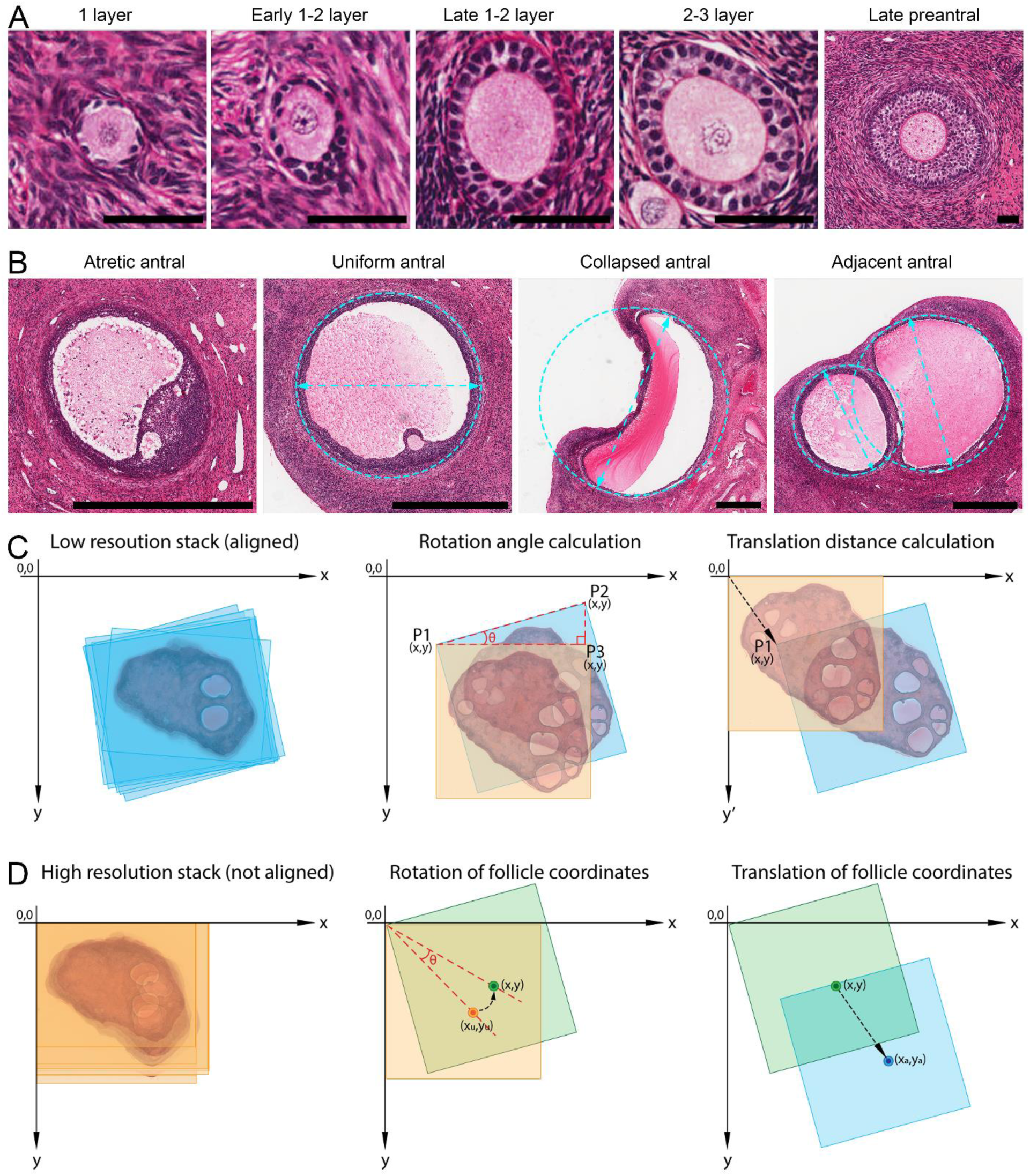
Follicle classification and determination of 3D coordinates. (A) Examples of follicles in different preantral stages of development based on the number of layers of granulosa cells. Late preantral follicles were recorded according to their diameter if they had more than 3 complete layers of granulosa cells (Scale bars = 50 µm). Follicle location was recorded as the x,y coordinate in the centre of the follicle. (B) Examples of antral follicles undergoing atresia or healthy follicles with uniform geometry, collapsed geometry caused by fixation or non-uniform geometry caused by follicles growing adjacent. Follicle diameters were determined at the widest point in the follicle and the midpoint was determined as the x,y coordinates at centre of the line traversing the widest diameter of the follicle. The z-coordinate was determined from the section number multiplied by 5 µm. (Scale bars = 1000 µm). (C) Image resolution of all histological images was reduced to enable a full ovary image series to be loaded and aligned in a Z-stack in FIJI software for obtaining large follicle coordinates. The amount of image rotation used in the alignment was determined using trigonometry based on the new x,y coordinates of the top left (P1) and top right (P2) corners of the image. Translation was defined as the P1 coordinate as the top-left corner of an unaligned image is at the origin (0,0). (D) Determination of the x,y coordinates of small follicles requires high resolution images that could not be loaded into a single Z-stack due to memory constraints. The coordinates of each small follicle were determined within individual images then the rotation and alignment steps were applied to all recorded x,y coordinates for small follicles. This step brings all small follicle coordinates in the database into alignment with the large follicle coordinates in 3D space.

**Supplemental figure 2.**
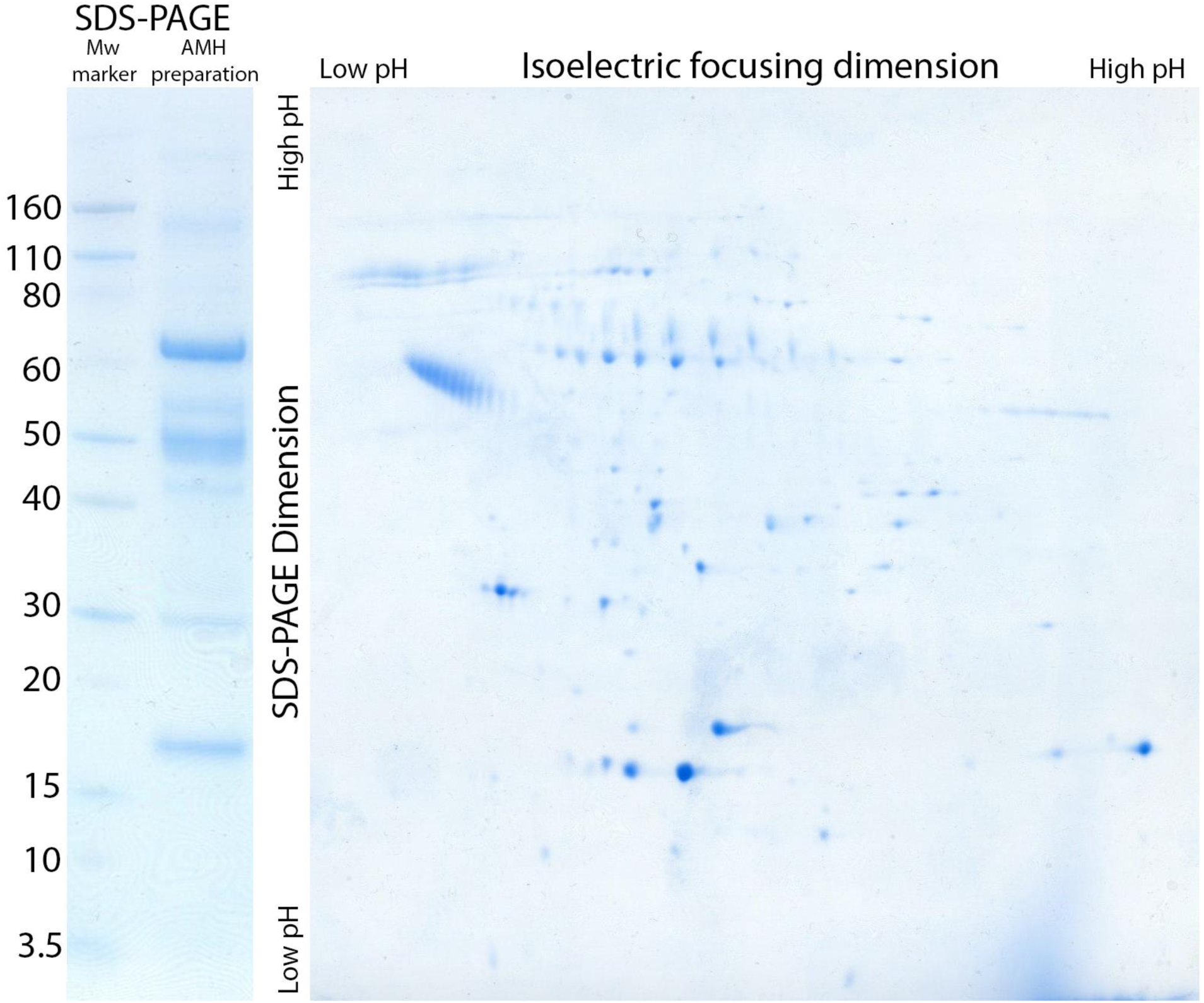
Large 2-dimensional blot of the AMH preparation (isoelectric focusing dimension horizontal, SDS-PAGE dimension vertical). The initial SDS-PAGE with molecular weight marker (kDa) is shown, left. The most prominent band at 72 kDa is the recombinant proAMH band.

**Supplemental figure 3.**
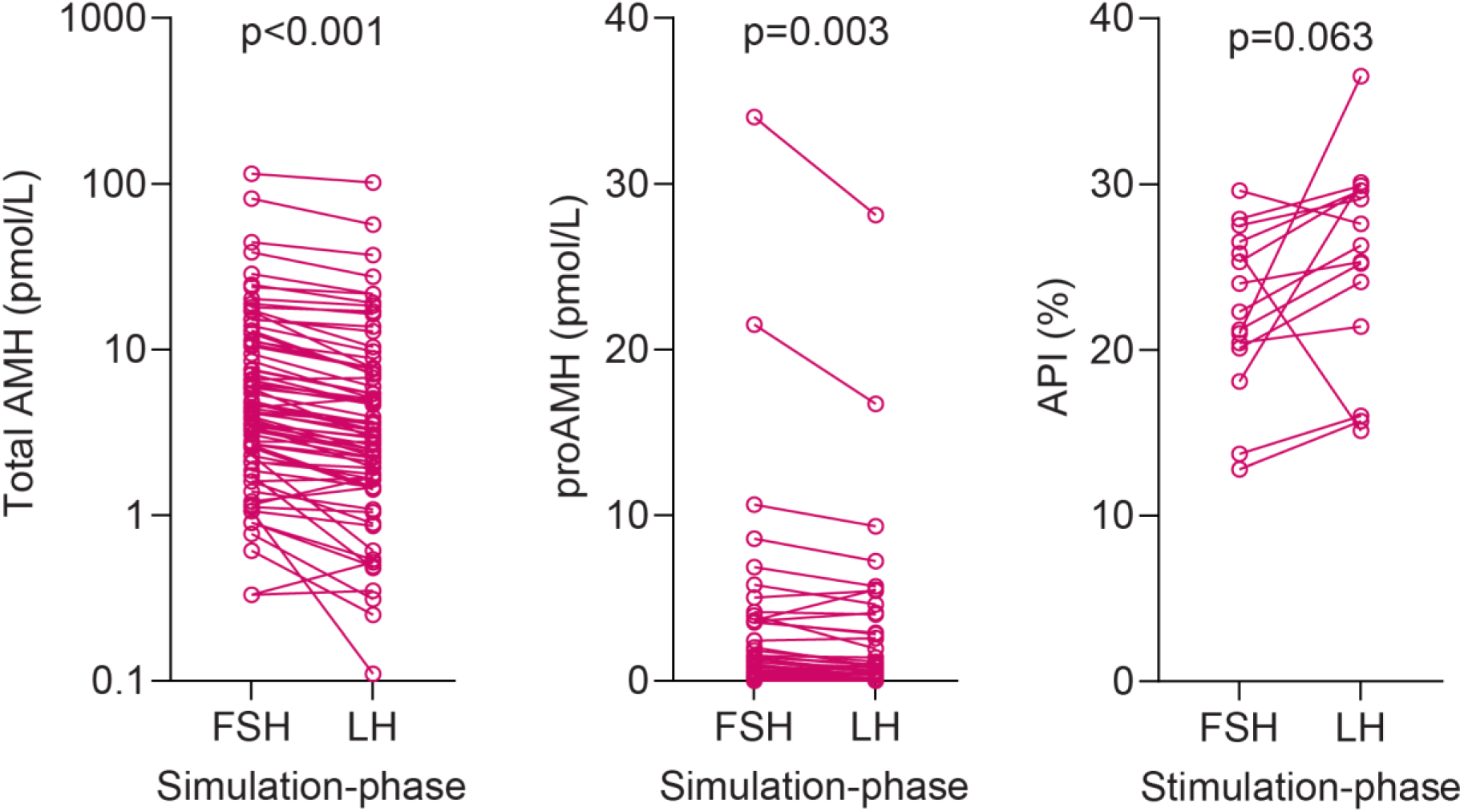
Serum total AMH, proAMH and the AMH prohormone index (API = [proAMH]/[total AMH] × 100) levels during controlled ovarian stimulation. Serum samples were obtained from patients undergoing controlled ovarian stimulation in both the FSH-stimulation phase and 24 hours after human chorionic gonadotropin injection (which stimulates the LH receptors on ovarian follicles). A total of 77 patients were assayed for total AMH and proAMH but only 15 patients had sufficiently high total AMH or proAMH concentrations for accurate calculation of API values. Paired, 2-tailed t-tests demonstrate significant decreased in total AMH and proAMH concentrations, but not the API, after LH-receptor stimulation.

**Supplemental figure 4.**
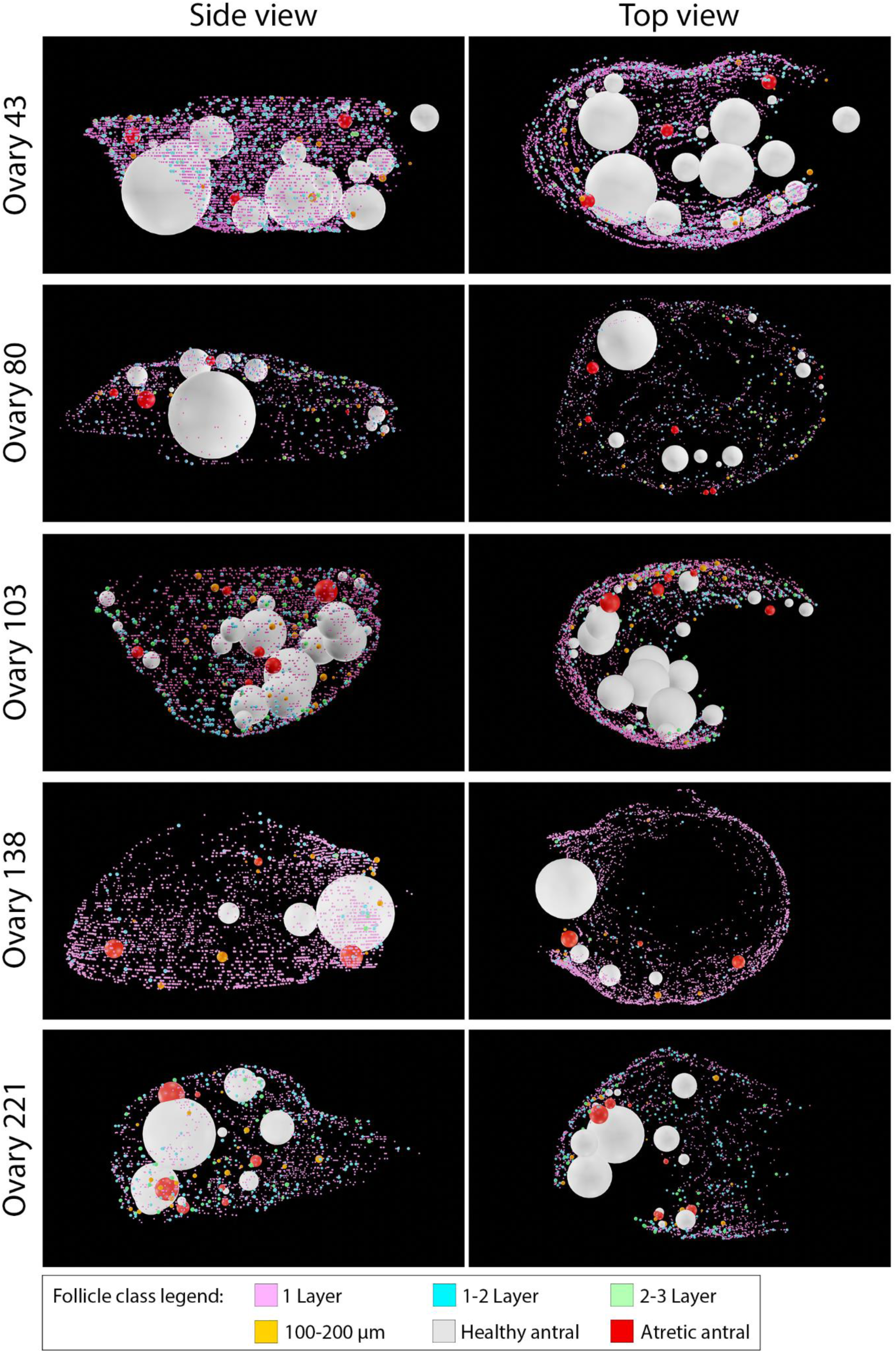
3D reconstructions of vehicle control-immunised sheep ovary follicle locations.

**Supplemental figure 5.**
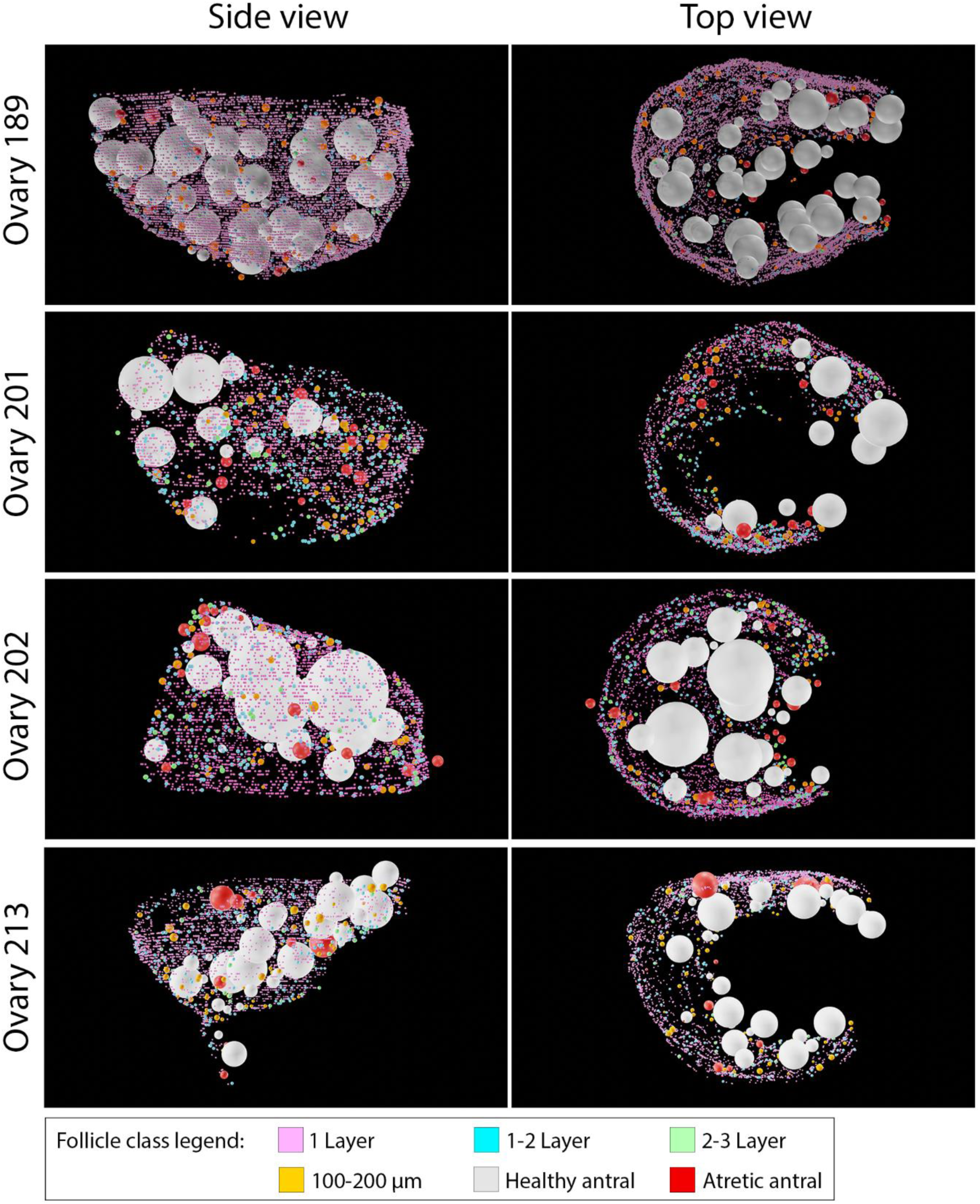
3D reconstructions of AMH-immunised sheep ovary follicle locations.

